# NeTOIF: A Network-based Approach for Time-Series Omics Data Imputation and Forecasting

**DOI:** 10.1101/2021.06.05.447209

**Authors:** Min Shi, Shamim Mollah

## Abstract

**Motivation:** High-throughput studies of biological systems are rapidly generating a wealth of ‘omics’-scale data. Many of these studies are time-series collecting proteomics and genomics data capturing dynamic observations. While time-series omics data are essential to unravel the mechanisms of various diseases, they often include missing (or incomplete) values resulting in data shortage. Data missing and shortage are especially problematic for downstream applications such as omics data integration and computational analyses that need complete and sufficient data representations. Data imputation and forecasting methods have been widely used to mitigate these issues. However, existing imputation and forecasting techniques typically address static omics data representing a single time point and perform forecasting on data with complete values. As a result, these techniques lack the ability to capture the time-ordered nature of data and cannot handle omics data containing missing values at multiple time points.

**Results:** We propose a network-based method for time-series omics data imputation and forecasting (NeTOIF) that handle omics data containing missing values at multiple time points. NeTOIF takes advantage of topological relationships (e.g., protein-protein and gene-gene interactions) among omics data samples and incorporates a graph convolutional network to first infer the missing values at different time points. Then, we combine these inferred values with the original omics data to perform time-series imputation and forecasting using a long short-term memory network. Evaluating NeTOIF with a proteomic and a genomic dataset demonstrated a distinct advantage of NeTOIF over existing data imputation and forecasting methods. The average mean square error of NeTOIF improved 11.3% for imputation and 6.4% for forcasting compared to the baseline methods.

## 1 Introduction

The development of high-throughput biology allows the production of large-scale omics datasets of varying levels ranging from genomics and epigenomics to transcriptomics and proteomics. In the past decades, the wide availability of various omics-scale data has greatly advanced our biological understanding of complex diseases such as breast cancer (Mollah and Subramaniam, 2020), neurodegenerative diseases (La Cognata *et al.*, 2021), and inflammatory bowel disease (Stylianou, 2018). Many of these advances have benefited from integrating multiple omics data making it possible to obtain comprehensive knowledge of the studied biological systems (Hasin *et al.*, 2017). For example, an integrative multi-omics analysis study of brains affected by Alzheimer’s disease has revealed associations of histones H3K27ac and H3K9ac in dysregulating transcription- and chromatin–gene feedback loops (Nativio *et al.*, 2020). In addition, an increasing number of computational studies, such as disease subtyping and biomarker prediction, have benefited from the integration of multiple omics data to achieve better classification and prediction accuracy (Subramanian *et al.*, 2020).

A common challenge of integrative and computational analyses is that omics data often contain missing values and limited observations, especially for sequential omics data observed from dynamic systems. These in turn hinder multi-omics data integration and many downstream analyses that require complete omics data representations. Missing values in the omics data may occur due to various experimental and technical reasons. For example, proteomics data show missing values due to the imperfect identification of coding sequences within a genome and the limited sensitivity of current peptide detection technologies (Albrecht *et al.*, 2010). A large portion of miRNAs is expressed below the detection limit in miRNA arrays, resulting in missing values in the output (Lin *et al.*, 2016). In addition to missing or incomplete values, shortage of data is another problem, especially for time-series omics data collected from dynamic observations. For example, capturing the dynamic interactions among growth promoting ligands, signaling proteins, histone modifications, and genes in the breast cancer microenvironment requires integrating multiple time-series omics data (e.g., proteomics and genomics) (Shi *et al.*, 2020). Typically these time-series datasets do not align perfectly, *i.e.,* each omic data only have observations in few time points. Therefore, it becomes challenging to incorporate multiple timeseries omics datasets and capture their biological dynamics. Omics data imputation and forecasting methods are therefore required to fill this gap.

Existing works on data imputation typically focus on static omics data as shown in the top panel of Fig.1a, including the Markov affinity-based graph imputation of cells (MAGIC) (van Dijk *et al.*, 2017), singular value decomposition imputation (SVDimpute) (Troyanskaya *et al.*, 2001), *k*-nearest neighbor imputation (KNNimpute) (Troyanskaya *et al.*, 2001), local least square imputation (LLS) (Kim *et al.*, 2005), and other methods. However, an obvious limitation of these methods is that they are inefficient at handling time-series omics data in which the time-ordered nature (e.g., time-series dependencies in data) of the data needs to be considered. To address this limitation, we propose here a novel network-based approach for time-series omics data imputation. As shown in Fig.1a, our method is able to model time-sequence information (e.g., time-series dependency relations between data representations) and allows for missing values at different time points while performing the target imputation. We additionally model the structure-dependency relations (topology) across data samples from their interaction networks (e.g., protein-protein or gene-gene interactions) to mitigate the influence of missing values at different time points and enhance the target imputation. Apart from the missing data imputation, the proposed method can also be used for omics data forecasting at the future time points. For the forecasting task, Fig.1b shows that our method allows the presence of missing values at different time points, unlike other existing methods that require input data representations to be complete. We evaluate the proposed method for timeseries imputation and forecasting using two real omics datasets: proteomics and genomics, and compare against five strong baseline methods in situations where the raw omics data contain different ratios of missing values. Comparative results show our method for time-series incomplete omics data imputation and forecasting outperforms other methods.

**Fig. 1:**
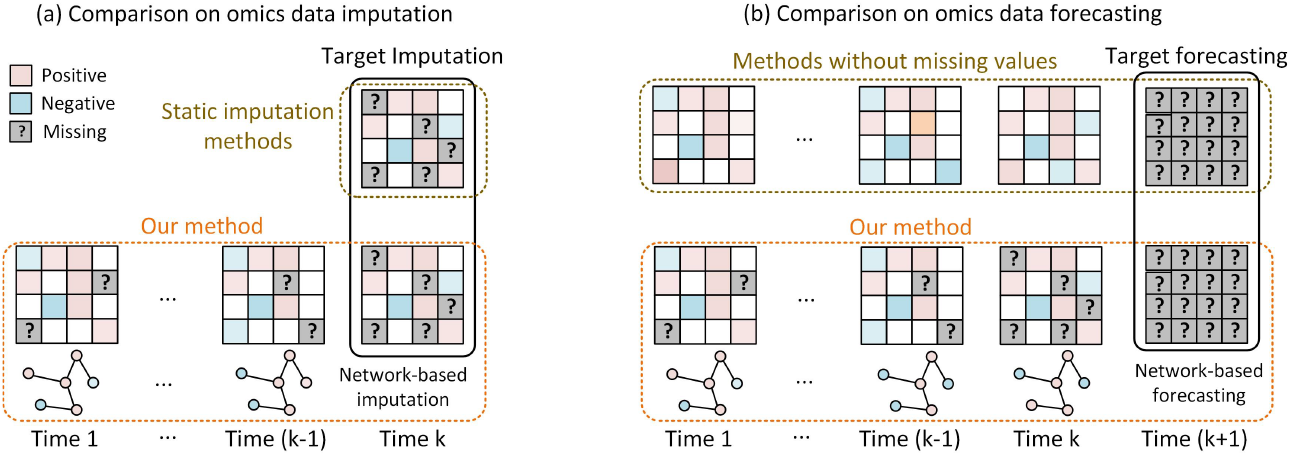
The differences between our method and existing methods with reference to data imputation and prediction. (a) Existing methods mainly focus on the static data imputation, whereas our method focuses on the time-series data imputation where missing values may happen at multiple time points. (b) For data forecasting, existing methods consider the observed data are complete, while our method allows missing values at multiple time points.

## 2 Materials and Methods

### 2.1 Problem Definition

Suppose the time-series omics data (e.g., *T* time points) be represented as 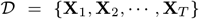, where **X**_*k*_ ∈ ℝ^*n*×*m*^ is the omics data representation (e.g., observations) corresponding to time point *k, n* denotes the number of data samples (e.g., proteins for proteomics data and genes for genomics data). *m* indicates the number of observations at multiple time points associated with each sample. In this paper, we focus on omics data where samples have structure dependencies such as proteinprotein and gene-gene interaction networks. Therefore, the omics data at each time point *k* can be represented as a network *G_k_* = {**V, E, X**_*k*_}, where **V** = {*v_i_*}_*i*=1,⋯,*n*_ is a set of samples and **E** = {*e_i,j_*}_*i,j*=1,⋯,*n*_ stores the structure relations among samples (e.g., edges). Note that omics data at different time points have the common sample node set **V** and edge set **E** but the data observations for nodes are varying, i.e., **X**_*k,i*_ ∈ ℝ^*m*^ only represents the observations for sample node *v_i_* at time point *k*.

#### 2.1.1 Time-series omics data imputation

The imputation task aims to impute the missing values involved in data representation 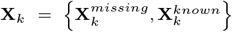 at a time point *k* ∈ [1, *T*], where 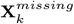 and 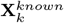 represent the sets of missing and known values respectively. We focus on time-series imputation where all previously observed data representations until time point *k* are used to train an imputation model, which then acts to impute the missing omics data at time point *k*. In addition, we allow the presence of missing values in omics data at different time points before *k* as shown in Fig.1a.

#### 2.1.2 Time-series omics data forecasting

As shown in Fig.1b, the forecasting task aims to predict the omics data representation at a future time point (*k* + 1) based on all the previous observations until time point *k*. Similar to the situation in data imputation, we allow the presence of missing values at different time points while forecasting the target omics data at time point (*k* + 1).

### 2.2 NeTOIF

The developed network-based approach for time-series omics data imputation and forecasting (NeTOIF) contains two major components: (1) a graph convolutional network (GCN) based structure dependency relationships modeling, and (2) a long short-term memory network (LSTM) based time-series dependency relationships modeling. The two components are trained in the unified framework (Fig.2).

**Fig. 2:**
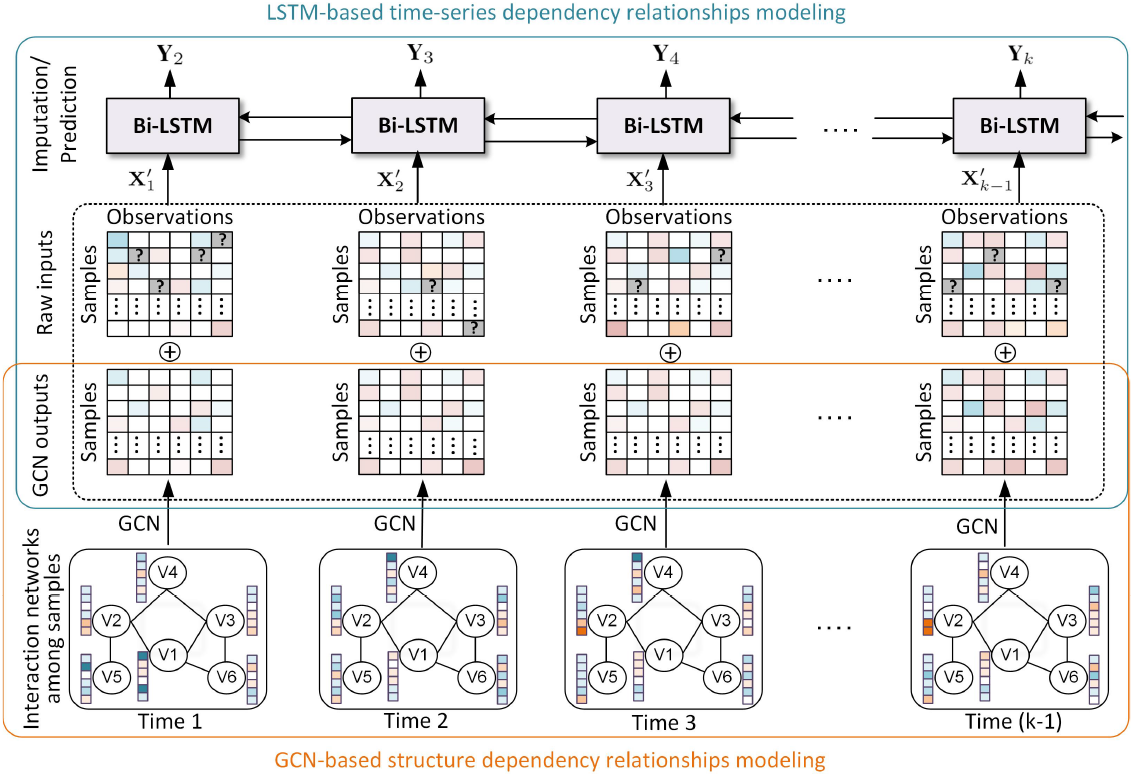
The framework of NeTOIF for both time-series omics data imputation and prediction. First, in the graph convolutional network (GCN) based structure dependency relationships modeling (**pink rectangle**), NeTOIF learns the missing values at different time points based on the omics sample interaction network. Then, NeTOIF model the time-series dependency relationship of omics data at different time points based on a long short-term memory (LSTM) network (**blue rectangle**). Both the GCN outputs and the original omics data at different time points are used as inputs of LSTM.

#### 2.2.1 GCN-based Structure Dependency Relationships Modeling

Our rationale for modeling the structure dependencies among omics data samples is twofold. First, network connections explicitly reveal rich functional relevance, such as physical pathway associations and orthology relations between samples, which can be used as reliable auxiliary information to strengthen the target data imputation and forecasting (Szklarczyk *et al.*, 2015). Moreover, structure relations can help infer the missing values in data representations at different time points, resulting to a more robust target imputation and forecasting. The second reason is based on the assumption that genes or proteins with neighborhood relationships on the network tend to have similar biological responses or observations. For instance, in a gene-gene co-expression network, all linked gene sample nodes should likely to have similar expression values (Van Dijk *et al.*, 2018). The missing values of genes at different time points can therefore be estimated from the known expression values of their linked genes.

Here, We adopted GCN (Kipf and Welling, 2016) to model the structure dependencies among samples (Fig.2), based on the principle that each sample node *v_i_* ∈ **V** at every time point *k* ∈ [1,*T*] generates its new observations (represented as a vector **X**_*k,i*_ ∈ ℝ^*m*^) by aggregating values from all immediate neighborhoods **N**(*v_i_*). Formally, with a single-layer GCN model, the new data representation of *v_i_* is calculated as:

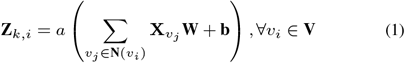

where **W** ∈ ℝ^*m*×*m*^ is a learnable parameter matrix and **b** is a learnable bias. *a* is a nonlinear activation function, *i.e., ReLU* (Nair and Hinton, 2010) represented by *f*(*x*) = max(0, *x*). The convolution representations calculated by Eq.1 with respect to all sample nodes at time point *k* can be entirely represented as (Kipf and Welling, 2016):

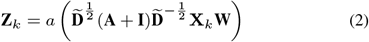

where **A** is the adjacency matrix of omics sample network, **I** is a samesize identity matrix and 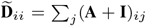. A single-layer GCN only captures the first-order structural dependency relationships (e.g., direct links) among omics data samples. We can easily stack more GCN layers to capture higher-order dependency relations between samples as:

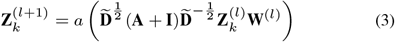

where 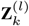 is the input data representations for the *l^th^* GCN layer as well as the output of the (*l* – 1)^*th*^ GCN layer, and **W**^(*l*)^ indicates the learnable weight parameter for the *l^th^* layer. The motivation to capture higher-order structure relationships with more GCN layers is that when the biological networks are complex (Dehmamy *et al.*, 2019), omics sample nodes that do not have direct links in the network may be correlated through their shared neighborhoods. For example, in the protein-protein interaction networks, proteins associated with the same signaling pathways tend to exhibit similar trend in expression but these pathway-based correlations may not be explicitly captured by the network structure (Wu *et al.*, 2010). In such a situation, we can capture the implicit high-order relationships between proteins using two or more GCN learning layers.

We used Eq.3 to obtain the GCN outputs at different time points until (*k* – 1) when performing the target data imputation and forecasting at time point k. In the following section, these GCN output data representations are combined with the original omics data representations for time-series imputation and forecasting using the bi-directional LSTM (Bi-LSTM) networks (Zhang *et al.*, 2015).

#### 2.2.2 LSTM-based Time-series Dependency Relationships Modeling

We adopted a Bi-LSTM network to model the time-series dependency relationships among omics samples (Fig.2). The NeTOIF model can be used for both omics data imputation and forecasting without any change to the model architecture.

The input for each time step of the Bi-LSTM network is a combination of the original omics data representations (e.g., may contain missing values) and the data representation output from the GCN network (through Eq.3). We adopted a pairwise addition (denoted by the symbol ⊕) of these two data representations at each time point *k* ∈ [1, *T*] as in Fig.2:

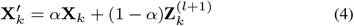

where *α* is a hyper parameter used to balance the importance between the original data representation **X**_*k*_ and the data representation 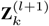 generated from the GCN. Then, the combined data representations undergo a series of non-linear transformation as in (Sahoo *et al.*, 2019):

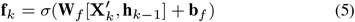

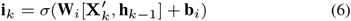

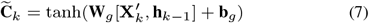

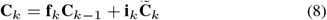

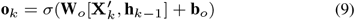

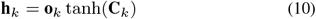

where **W**_*f*_, **W**_*i*_, **W**_*g*_ and **W**_*o*_ are trainable weight parameters for the LSTM networks, and **b**_*f*_, **b**_*i*_, **b**_*g*_ and **b**_*o*_ are the respective trainable bias parameters. *σ* is an activation function referred to as the sigmoid transformation (Han and Moraga, 1995), i.e., 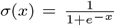. Finally, the bidirectional LSTM outputs 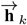 and 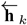 from the temporal and reverse orders are added to represent the final output as:

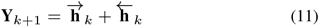

The Bi-LSTM output by Eq.11 allows the capturing of the time-series dependency relationships between adjacent data representations from the two opposite directions shown in Fig.2.

#### 2.2.3 Training NeTOIF for Omics Data Imputation and Forecasting

NeTOIF is a unified model but the schemes to construct training samples for data imputation and forecasting are different. For data imputation, all previous data representations before time *k* (e.g., fixed window) are used as inputs of NeTOIF to impute the missing values in the data representation at time point *k* ∈ [1,*T*] (Fig.3a). Model weight parameters are trained and optimized by minimizing the mean square error (MSE) between the imputed values and the respective actual values as:

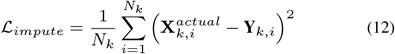

where 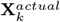 stores all the actual values and *N_k_* is number of actual values at time *k*. **Y**_*k*_ is obtained by Eq.11.

**Fig. 3:**
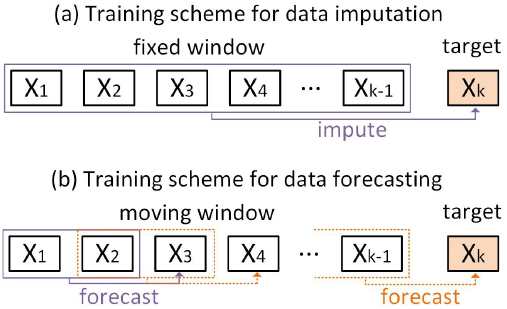
Training schemes of NeTOIF. (a) For data imputation, all previous data before time *k* are used to impute missing values in the data at time point *k*. (b) For data forecasting, a moving window slides along data at different time points, where all data within each window are used to forecast the data at a future time point.

In comparison, for data forecasting in Fig.3b, a moving window (of size *s*) is defined to construct training samples, where all data representations within each window context are used to forecast the data representations at a future time point. All data before time *k* are used to train the model and minimize the following MSE loss as:

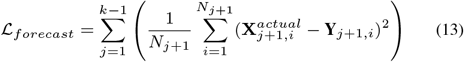

where 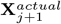 stores all actual values at the future time point (*j* + 1) and **Y**_*j*+1_ is computed by Eq.11.

We then optimized Eq.12 and Eq.13 using *Adam*, a stochastic gradient-based optimization method (Kingma and Ba, 2014). The detailed procedures for omics data imputation and forecasting based NeTOIF are respectively summarized in Algorithm 1 and Algorithm 2, where **W**\* represents any learnable weight parameter involved in NeTOIF.

### 2.3 Time-Series Omics Datasets and Preprocessing

We used a time-series proteomics and a time-series genomics dataset to evaluate the proposed approach as follows.

#### 2.3.1 Reverse Phase Protein Array Data

The reverse phase protein array (RPPA) proteomics data were downloaded from the Synapse platform (https://www.synapse.org/#!Synapse:syn12555331). The dataset consists of highly sensitive and selective antibody-based measurements of 295 proteins and phosphoproteins in breast epithelial cells after individual treatments with six different growth ligands: epidermal growth factor (EGF), hepatocyte growth factor (HGF), oncostatin M (OSM), bone morphogenetic protein 2 (BMP2), transforming growth factor beta (TGFB), and interferon gamma-1b (IFNG). Each data sample contains three replicates for each of these ligands measured at five time points (1, 4, 8, 24, and 48 hours after treated with ligands).

**Algorithm 1.**
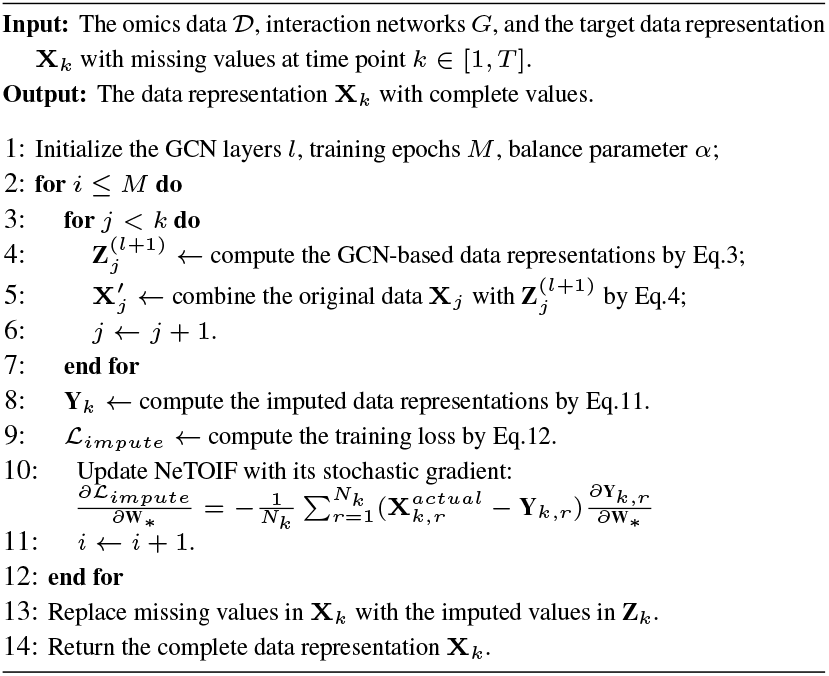
Time-series omics data imputation based on NeTOIF

**Algorithm 2.**
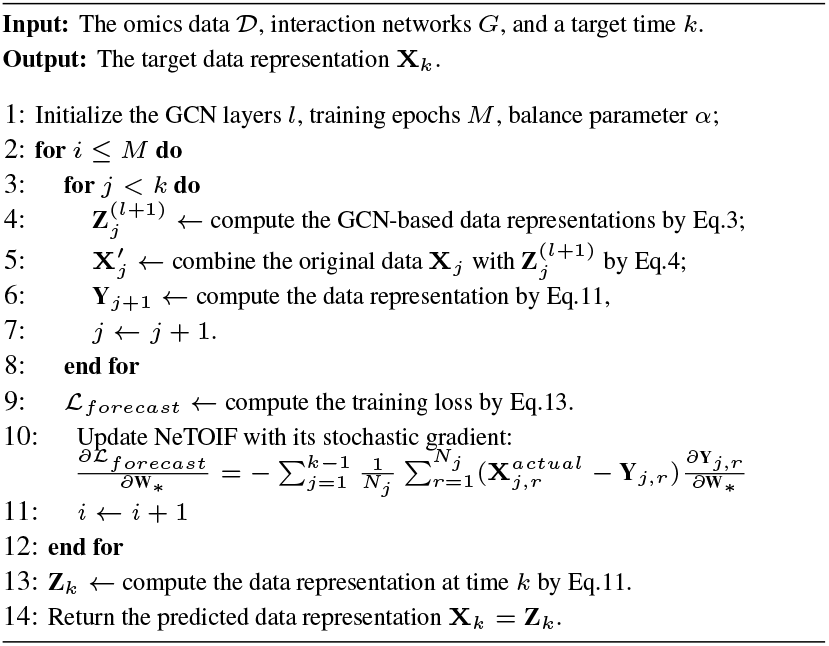
Time-series omics data forecasting based on NeTOIF

The three replicates for each sample were averaged to obtain a single data representation at each of the five time points. A protein-protein interaction (PPI) network was constructed from the STRING database (https://string-db.org/), which contains 989 links among the 295 proteins.

#### 2.3.2 Genome-Wide Gene Expression Data

The genome-wide gene expression (GE) data were published in literature (Mutarelli *et al.*, 2008). Human estrogen-responsive breast cancer cells (ZR-75.1) cultured in steroid-free medium for 4 days were stimulated with a mitogenic dose (10nM) of 17*β*-estradiol and RNA was extracted before hormonal stimulation or after 1, 2, 4, 6, 8, 12, 16, 20, 24, 28 or 32 hours of hormonal stimulation.

The gene expression data for the 4-, 8-, 12-, 16-, 20-, 24- and 32-hour time points were retained to form a time-series (e.g., equal intervals) genomics dataset in which adjacent time points have an identical interval of 4 hours. Similarly, a gene-gene interaction network was constructed from the STRING database (https://string-db.org/), which contains 644 genes and 4584 links among them.

## 3 Experiments

### 3.1 Compared Methods

We compare the developed NeTOIF with five baseline methods used for either static or time-series data imputation and forecasting as follows:

- **MAGIC (MAGIC)** (van Dijk *et al.*, 2017): MAGIC is a graph-based method originally designed for imputing missing values of the static single-cell gene expression data. The main idea of MAGIC is to share information across similar cells and fill in missing gene expression values of a given cell based on the neighborhood cells. This method is only used for static data imputation at a single time point.
- **Bayesian Bridge Regression (Bayesian)** (M Mostafa *et al.*, 2020): This is a regression model with an additional regularization parameter for the coefficients, where the prior for the coefficient is given by a spherical Gaussian. This method adopts the same training scheme as our method for time-series imputation and forecasting shown in Fig.3.
- **Support Vector Regression (SVR)** (Wang *et al.*, 2006): This is extended from the support vector machine model for regression problems. In SVR the inputs are mapped into a higher dimensional space in a non-linear manner based on the kernel function. This method uses the same imputation and forecasting schemes shown in Fig.3.
- **Long Short-Term Memory Networks (LSTM)** (Hochreiter and Schmidhuber, 1997): This is a neural network model that requires timeseries data as inputs. LSTM is able to model the long-time dependency relations between temporal data samples and has been widely used for time-series regression and prediction tasks. This method uses the same imputation and forecasting schemes shown in Fig.3.
- **Convolutional Neural Networks (CNN)** (Aghdam and Heravi, 2017): This is a neural network model widely used for image and text processing. The hidden layers of a CNN typically consist of a series of convolutional layers that convolve with a multiplication or other dot product. This method uses the same imputation and forecasting schemes shown in Fig.3.

Among all compared methods in the experiment, NeTOIF, Bayesian, SVR, LSTM and CNN are methods which are able to capture the timeseries dependency relationships between omics samples, whereas MAGIC can only model the static relations between samples at a single time point.

### 3.2 Evaluation Metrics

We adopted Mean Square Error (MSE) and Mean Absolute Error (MAE) to quantify the performance of all baseline methods and the proposed method in this paper. The MAE and MSE are defined as follows:

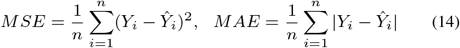

where *n* is the number of data samples, and *Y_i_* and 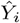 are the actual and imputed (forecasted) values for the *i^th^* data sample. Smaller MSE and MAE values indicate better imputation or forecasting results.

### 3.3 Experimental Settings

We adopted a single-layer GCN in NeTOIF to learn the omics sample networks in both imputation and forecasting tasks. We used *Adam* (Kingma and Ba, 2014) to train NeTOIF with a learning rate equals to 5-e3. We perform extensive experiments to test the sensitivity of the balance parameter *α* and the window size *s* used in the forecasting task. In the data imputation task, we used parameter *ρ* to represent the ratio of missing values in the original data representations at a given time point *k*. In addition, we used parameter λ to represent the ratio of missing values in each of previous data representations before *k* while performing the target data imputation or forecasting. We ran the experiments 20 times and reported the average results and their standard deviations. We used a pairwise Student’s t-test to determine if the results of a given pair of methods are statistically different from each other. Results of NeTOIF were considered to reach significance at *p* ≤0.05 and are indicated with asterisks (*, *p* ≤ 0.05; **, *p* ≤ 0.01; ***, *p* ≤ 0.001).

In this paper, we evaluated NeTOIF for time-series data imputation on both RPPA and GE datasets. Since the time intervals between different time points in RPPA are not equivalent, we only used GE data to test the time-series data forecasting performance of NeTOIF.

### 3.4 Result and Discussion

The omics data imputation and forecasting results are first presented and discussed in this section. Then, extensive experiments are designed to test the sensitivities of the balance parameter *α* and window size *s*.

#### 3.4.1 The omics data imputation results

From the imputation results on the RPPA data using different ratios of missing values at the 48 hour time point (Fig.4), we observe that the NeTOIF, CNN, LSTM, SVR and Bayesian methods significantly (***) outperform MAGIC in both MSE and MAE. All methods except MAGIC have utilized the time-series dependency relationships between samples, which are beneficial information to improve the missing value imputation. Changes in the omics data over time can be seen as a function of the time variable, meaning that all omics data prior to the 48 hour time point can be used to impute missing values at 48-hour time point. The developed method NeTOIF performs generally better than other baselines (** when *ρ* = [0.3, 0.5] for MSE, and * when *ρ* = 0.3 for MAE). For example, when *ρ* = 0.3 the MSE result of NeTOIF improved 57.4% compared to MAGIC, 29.7% compared to Bayesian, 29.3% compared to SVR, 32.2% compared to LSTM, and 29.6% compared to CNN, respectively. This is because a smaller value of *ρ* means that more observed data are available for training the weight parameters in NeTOIF, thereby producing a better-performing imputation model.

**Fig. 4:**
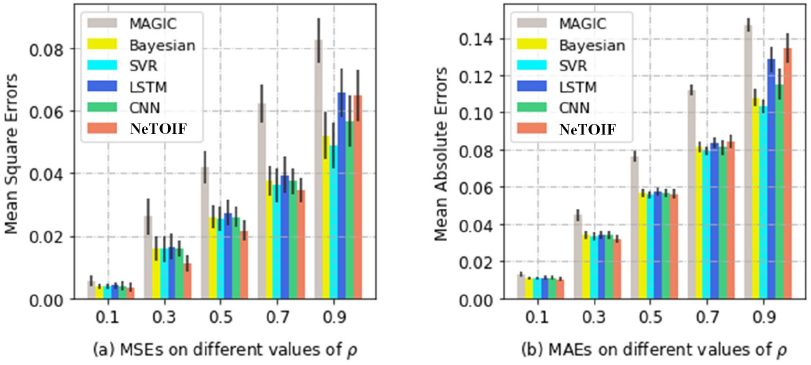
Comparison of imputation results on the RPPA data at 48-hour time point. The x-axes represent the different ratios of missing values in the original data representation controlled by *ρ*. The y-axes represent the MSE (a) or MAE (b) values.

As discussed previously, while performing the time-series imputation, NeTOIF allows the presence of missing values at multiple time points. This is shown when different ratios (λ) of actual values were randomly removed and replaced with zeros at each previous time point before the target time point under imputation (Fig.5). The MSE and MAE of all timeseries based methods gradually increase with the increase of λ (Fig.5). This is because the time-series information becomes unreliable when the data contain a high level of missing values. With different values of λ, we observe that NeTOIF performs generally better than all other baseline methods regarding the MSE results (Fig.5a, ** when λ ≤ 0.7 for MSE), but in contrast, the MAE result is similar to other time-series based baseline methods (Fig.5b). Similarly, Fig.6 shows the comparison results (** when λ = 0.3 and *** when λ > 0.3) on the GE data, NeTOIF performs generally better than baseline methods regarding both MSE and MAE results here. In contrast, the MSE and the MAE stay highly independent of the variation of λ using the method MAGIC (Fig.5 and Fig.6). This is because MAGIC does not consider the time-series information and is not affected by missing values at previous time points before the target time point under imputation.

**Fig. 5:**
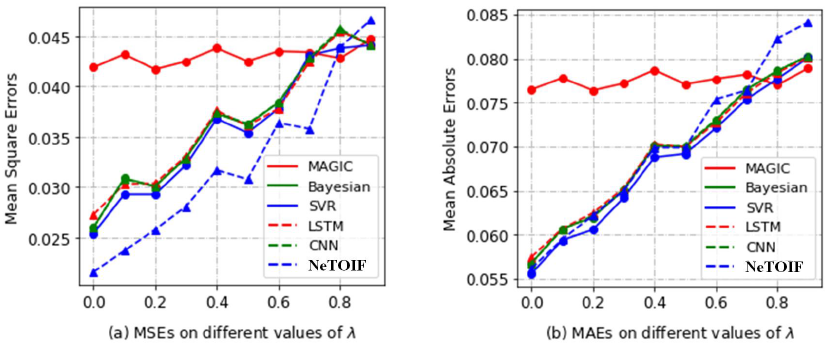
Comparison of imputation results on the RPPA data at the 48-hour time point. The x-axes represent different ratios of missing values at 1-, 4-, 8- and 24-hour time points controlled by λ. The y-axes represent the MSE (a) or MAE (b) values.

**Fig. 6:**
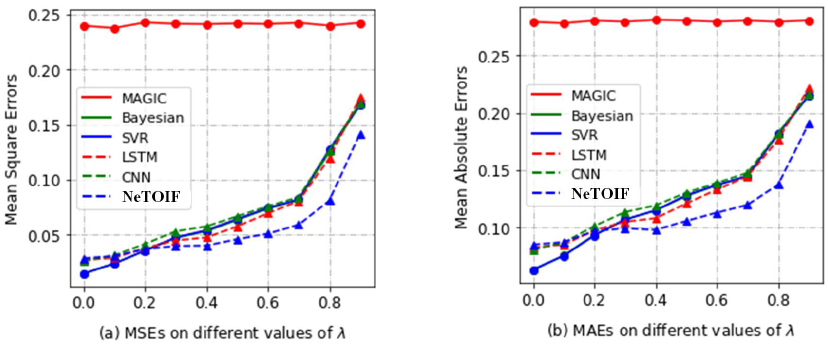
Comparison of imputation results on the GE data at the 32-hour time point. The x-axes represent different ratios of missing values at 4-, 8-, 12-, 16-, 20-, 24- and 28-hour time points controlled by λ. The y-axes represent the MSE (a) or MAE (b) values.

Fig.7 shows the imputed RPPA (48-hour time point) and GE data representations (32-hour time point) compared with the respective actual data representations. We observe that NeTOIF has produced promising imputation results on both datasets. For example, for the GE data the imputed directions (blue indicates negative and red indicates positive) of gene expressions are generally aligned with the actual data (Fig.7b). Fig.8 shows the impact of balance parameter *α* used in Eq.4. We can see that the optimal value setting for α tends to be smaller when λ is larger. A smaller value of *α* indicates that topological relationship information plays a more important role in the imputation. The above phenomenon demonstrates that for time-series proteomics data that contain a high ratio of missing values at different time points, the topological relationships between samples are helpful to improve the imputation accuracy.

**Fig. 7:**
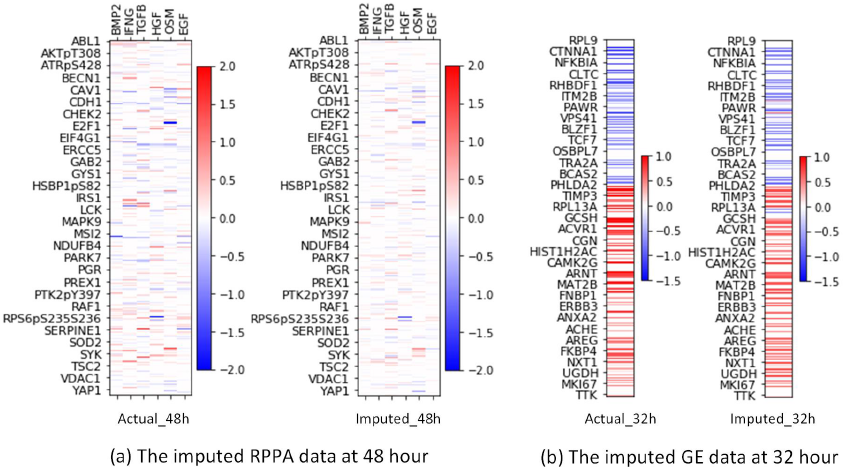
The imputed RPPA (a) and GE data (b) by NeTOIF compared with the respective actual data representations (*ρ* = 0.5).

**Fig. 8:**
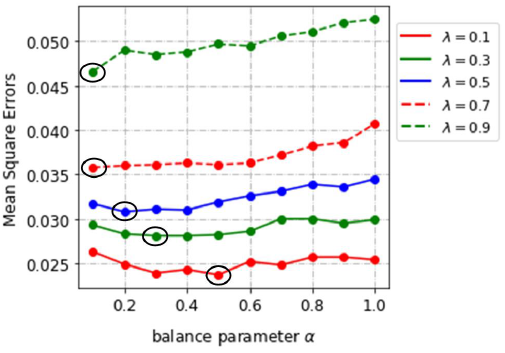
The influence of the balance parameter *α* on RPPA data in the imputation task. The best values of *α* for different settings of λ are circled.

#### 3.4.2 The omics data forecasting results

Table 1 compares the forecasting performance of various methods on the GE data at the 32-hour time point. NeTOIF significantly outperforms all baseline methods regarding MSE and MAE results, though all methods declined with the increase of λ. For example, the average MSE of NeTOIF improved by 6.4% compared to the strongest baseline method LSTM. Fig.9 shows that the forecasted GE data at the 28- and 32-hour time points are comparable with the actual data. The value distributions of gene expressions in the forecasted data are well aligned with the actual data. In the forecasting task, we use all time points within a window to forecast the next time point in the future. Larger window size (*s*) tends to deliver better MSE results (Fig.10), because it allows to observe more sufficient changing patterns of samples with time, which help to predict trends at future time points.

**Fig. 9:**
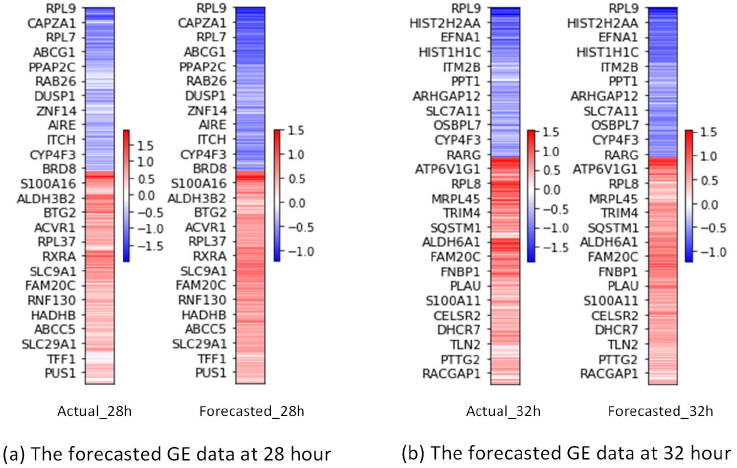
The forecasted GE data by NeTOIF compared with the respective actual data representations.

**Fig. 10:**
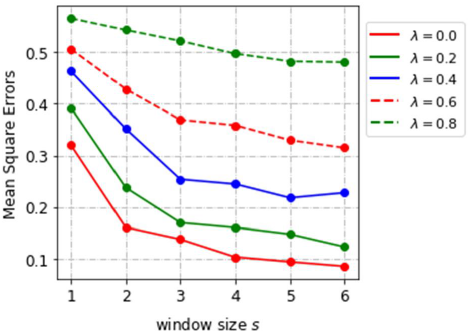
The influence of the window size *s* on GE data forecasting at 32 hour-time point with regards to different values of λ.

**Table 1.** Comparison of the forecasting results on the GE data at 32 hour w.r.t different ratios of missing values in the historical time points controlled by λ. The best results are highlighted and marked with asterisks (*, *p* ≤ 0.05; **, *p* ≤ 0.01; ***, *p* ≤ 0.001) if they are significantly different from others.

| Metrics | Mean Square Error |  |  |  |  | Mean Absolute Error |  |  |  |  |
| --- | --- | --- | --- | --- | --- | --- | --- | --- | --- | --- |
| $\lambda$ | 0.0 | 0.2 | 0.4 | 0.6 | 0.8 | 0.0 | 0.2 | 0.4 | 0.6 | 0.8 |
| Bayesian | 0.1089 | 0.1875 | 0.2859 | 0.3987 | 0.5197 | 0.2537 | 0.3392 | 0.4339 | 0.5404 | 0.6413 |
| SVR | 0.1075 | 0.1769 | 0.2987 | 0.4810 | 0.5457 | 0.2507 | 0.3257 | 0.4464 | 0.6116 | 0.6590 |
| LSTM | 0.1120 | 0.1704 | 0.2687 | 0.3775 | 0.5120 | 0.2566 | 0.3208 | 0.4141 | 0.5192 | 0.6347 |
| CNN | 0.1084 | 0.1883 | 0.2799 | 0.4005 | 0.5200 | 0.2529 | 0.3403 | 0.4275 | 0.5421 | 0.6414 |
| NeTOIF | **0.1033\*\*\***<br>($\pm 0.0030$) | **0.1511\*\*\***<br>($\pm 0.0049$) | **0.2445\*\*\***<br>($\pm 0.0054$) | **0.3523\*\*\***<br>($\pm 0.0044$) | **0.4964\*\*\***<br>($\pm 0.0024$) | **0.2449\*\*\***<br>($\pm 0.0036$) | **0.3040\*\*\***<br>($\pm 0.0065$) | **0.3966\*\*\***<br>($\pm 0.0062$) | **0.5091\*\*\***<br>($\pm 0.0027$) | **0.6212\*\*\***<br>($\pm 0.0022$) |

## 4 Conclusion

Missing values are common in time-series omics data such as proteomics and genomics data, which disrupt many downstream analyses and applications. This further complicates analyses of time-series data which not only contain missing values at multiple time points but also present a shortage of data observations (e.g., omics data at more time points are desired). In this paper, we proposed a novel network-based method called NeTOIF that incorporates both structural and time-series dependency relationships between omics samples to perform data imputation and forecasting. For our study, we used proteomics (RPPA) and genomics (GE) time-series datasets to validate the utility of the NeTOIF against five baseline methods. Our NeTOIF generally outperformed other baseline methods for time-series omics data imputation and forecasting, where we believe the performance gain was achieved by incorporating topological structures of genes and proteins interactions.

In this study, we mainly focused on time-series omics data where a network of interactions was available for capturing the topological relationships between samples. In the future, we plan to extend our method by automatically learning the topology structures between the omics samples and develop efficient models that can handle omics data without topology dependency.

## Contributions

Original concept: S.M., M.S.; experimental design: M.S., S.M.; data analysis: M.S.; implementation: M.S., S.M.; data interpretation: M.S., S.M.; manuscript writing: M.S., S.M.

## Acknowledgements

We want to thank NIH LINCS consortium for making RPPA data available.

